# Comparative cardiotoxicity assessment of bisphenol chemicals and estradiol using human induced pluripotent stem cell-derived cardiomyocytes

**DOI:** 10.1101/2023.09.13.557564

**Authors:** Blake L. Cooper, Shatha Salameh, Nikki Gillum Posnack

**Author notes:** **Corresponding author:** Nikki Gillum Posnack, PhD, Sheikh Zayed Institute for Pediatric Surgical Innovation, 111 Michigan Avenue, NW, Washington DC 20010.

## Abstract

**Background:** Bisphenol A (BPA) is commonly used to manufacture consumer and medical-grade plastics. Due to health concerns, BPA substitutes are being incorporated – including bisphenol S (BPS) and bisphenol F (BPF) – without a comprehensive understanding of their toxicological profile.

**Objective:** Previous studies suggest that bisphenol chemicals perturb cardiac electrophysiology in a manner that is similar to 17β-estradiol (E2). We aimed to compare the effects of E2 with BPA, BPF, and BPS using human induced pluripotent stem cell-derived cardiomyocytes (hiPSC-CM).

**Methods:** Cardiac parameters were evaluated using microelectrode array (MEA) technology and live-cell fluorescent imaging at baseline and in response to chemical exposure (0.001-100 μM).

**Results:** Cardiac metrics remained relatively stable after exposure to nanomolar concentrations (1-1,000 nM) of E2, BPA, BPF, or BPS. At higher micromolar concentrations, chemical exposures resulted in a decrease in the depolarizing spike amplitude, shorter field potential and action potential duration, shorter calcium transient duration, and decrease in hiPSC-CM contractility (E2 ≥ BPA ≥ BPF >> BPS). Cardiomyocyte physiology was largely undisturbed by BPS exposure. BPA-induced effects were exaggerated when co-administered with an L-type calcium channel antagonist (verapamil) or E2 - and reduced when co-administered with an L-type calcium channel agonist (Bay K8644) or an estrogen receptor alpha antagonist (MPP). E2-induced effects generally mirrored those of BPA, but were not exaggerated by co-administration with an L-type calcium channel antagonist.

**Discussion:** Collectively across multiple cardiac endpoints, E2 was the most potent and BPS was the least potent disruptor of hiPSC-CM function. Although the observed cardiac effects of E2 and BPA were similar, a few distinct differences suggest that these chemicals may act (in part) through different mechanisms. hiPSC-CM are a useful model for screening cardiotoxic chemicals, nevertheless, the described *in vitro* findings should be validated using a more complex *ex vivo* and/or *in vivo* model.

## INTRODUCTION

Bisphenol A (BPA) is a high production volume chemical that is widely used in the manufacturing of epoxy resins and consumer and medical-grade polycarbonate plastics (Duty et al. 2013; Iribarne-Durán et al. 2019; Vandenberg et al. 2007). Given its relative abundance in the environment, biomonitoring studies indicate that >90% of the United States population is routinely exposed to BPA (Calafat et al. 2005, 2008; Lehmler et al. 2018; Vandenberg et al. 2010; Ye et al. 2015). Of concern, heightened BPA exposure is associated with adverse cardiovascular outcomes (Lind et al. 2021; Ramadan et al. 2020), including an increased risk of hypertension, atherosclerosis, coronary heart disease, and depressed heart rate variability (Bae et al. 2012; Bae and Hong 2015; Melzer et al. 2010; Moon et al. 2021; Shankar and Teppala 2012). Further, longitudinal cohort studies performed in the United States suggest that individuals with elevated BPA exposure have a ∼40% higher risk of cardiovascular and all-cause mortality (Bao et al. 2020; Chen et al. 2023). These health associations have prompted a renewed interest in understanding the link between bisphenol chemical exposure and cardiovascular toxicity.

To date, experimental studies suggest that the biological effects of BPA may be mediated through estrogen receptor signaling and/or inhibition of cardiac ion channels (Cooper and Posnack 2022). Multiple groups have demonstrated that BPA rapidly and reversibly blocks voltage-gated sodium and calcium channels in a dose-dependent manner (Deutschmann et al. 2013; Feiteiro et al. 2018; Hyun et al. 2021; Liang et al. 2014; Michaela et al. 2014; O’Reilly et al. 2012; Prudencio et al. 2021; Wang et al. 2011). In cardiac tissue, sodium channel blockers slow depolarization and conduction velocity - while calcium channel blockers reduce heart rate and slow atrioventricular conduction. Calcium flux also plays a vital role in cardiomyocyte excitation-contraction coupling, which is also disrupted by BPA exposure (Hyun et al. 2021; Posnack et al. 2015; Ramadan et al. 2018; Yan et al. 2011). A few studies have also demonstrated that BPA modulates intracellular calcium handling through estrogen receptor signaling – akin to 17β-estradiol (Belcher et al. 2011; Yan et al. 2011, 2013). *In vitro* studies report an immediate decrease in peak calcium and potassium current upon exposure to 17β-estradiol (E2), the predominant circulating estrogen in women (Berger et al. 1997; Jiang et al. 1992; Meyer et al. 1998).

Over time, BPA-related health concerns have shifted consumer preference for non-BPA products and influenced stricter manufacturing regulations, which in turn, has led to the introduction of new replacement chemicals – many of which structurally resemble BPA (Lehmler et al. 2018; Yu et al. 2015). Unfortunately, the safety profile of these alternative chemicals remains unclear, with some studies suggesting that these “regrettable substitutions” have comparable biological effects to BPA (Delfosse et al. 2012; Kojima et al. 2019; Trasande 2017; Wang et al. 2022). For example, one study found that bisphenol S (BPS) altered intracellular calcium signaling and increased the incidence of arrhythmias in a manner comparable to BPA. While another study reported that BPS is a less potent antagonist of L-type calcium channels with negligible effects on cardiac electrophysiology (Prudencio et al. 2021), which could be attributed to slight differences in its chemical structure (Deutschmann et al. 2013).

In the presented study, we aimed to evaluate the direct effects of BPA on cardiac electrophysiology and compare its potency to both E2 and alternative bisphenol chemicals (e.g., bisphenol S (BPS) and bisphenol F (BPF)). Based on prior work identifying BPA as a potent inhibitor of L-type calcium channels, we anticipated that BPA would perturb cardiac electrophysiology and calcium handling to a greater extent than BPS. To date, bisphenol toxicology studies have largely been limited to rodent model systems, which requires the careful consideration of species-specific differences when predicting human risk (Edwards and Louch 2017; Ripplinger et al. 2022). With this in mind, we evaluated the cardiotoxicity of bisphenol chemicals using human induced pluripotent stem cell-derived cardiomyocytes (hiPSC-CM). hiPSC-CM are considered a promising “new approach methodology” for cardiac risk assessment based on their similarity to human cardiomyocyte ion channel expression, action potential morphology, and contractile machinery (Chen et al. 2016; Gintant et al. 2017; Parish et al. 2020; Sinnecker et al. 2013).

## MATERIALS AND METHODS

### Reagents

A list of all chemical reagents is provided in **Supplemental Table 1**. Stock solutions were prepared in >99.9% anhydrous dimethyl sulfoxide (DMSO) and working concentrations were diluted in cell culture media. Treatment groups included: 0.01–100 μM E2, BPA, BPF, or BPS; 10 nM Bay K8644 (BayK); 100 nM verapamil; 10 μM 4,4’,4’’-(4-propyl-[1H]-pyrazole-1,3,5-triyl)trisphenol (PPT); 10 μM diarylpropionitrile (DPN); 1 μM ICI 182,780 (ICI); 1 μM 1,3-bis(4-hydroxyphenyl)-4-methyl-5-[4-(2-piperidinylethoxy)phenol]-1H-pyrazole dihydrochloride (MPP); 1 μM 4-[2-Phenyl-5,7-bis(trifluoromethyl)pyrazolo[1,5-a]pyrimidin-3-yl]phenol (PHTPP); 1 μM progesterone. The range of bisphenol concentrations was selected to mimic environmental exposure (1–100 nM), maximum clinical or occupational exposure (1-10 μM), and supraphysiological exposure levels (>10 μM) (Bousoumah et al. 2021; Ramadan et al. 2020). Pharmacological agent concentrations were selected based on previous reports (Hamilton et al. 1987; Long et al. 1998; Paterni et al. 2014; Zhou et al. 2009) and preliminary potency experiments. For drug combinations, hiPSC-CM were pretreated with a pharmacological agent (15-20 min), then BPA or E2 was added (15-20 min) before recording electrical signals. DMSO was used as a vehicle control (0.1%). To reduce the risk of environmental contamination, non-polycarbonate bisphenol-free plastic consumables were used for experiments (including polystyrene cell culture plates from Axion BioSystems).

### hiPSC-CM preparation

hiPSC-CM (iCell cardiomyocytes^2^, donor #01434, Fujifilm Cellular Dynamics) were thawed and plated on fibronectin-coated glass-bottom dishes (100,000 cells/cm^2^) or microelectrode array plates (40,000 cells/well; 24-well plate, Axion Biosystems), as previously described (Cooper et al. 2021; Prudencio et al. 2021). Female donor hiPSC-CM were used, as prior studies suggest that bisphenol toxicity may be exaggerated in female cells (Belcher et al. 2011; Gao et al. 2015). Cells were maintained under standard cell culture conditions (37°C, 5% CO_2_) in iCell cardiomyocyte maintenance media (Fujifilm Cellular Dynamics) supplemented with 1 mg/mL ciprofloxacin (Sigma Aldrich). Four days after plating, hiPSC-CM formed a confluent monolayer and were subsequently used for ongoing experiments.

### Microelectrode array (MEA) recordings

MEA plates were equilibrated for 15-20 min in a temperature-controlled cell monitoring system (Maestro Edge system, Axion Biosystems); electrical signals were recorded at baseline and after chemical treatment (**Figure 1A-M**). Electrical signals were recorded during spontaneous rhythm and in response to external stimulation (1-2 Hz). Cardiac parameters included: spontaneous beating rate, spike amplitude, conduction velocity, extracellular field potential duration rate-corrected with the Fridericia formula (FPDc), action potential duration (APD), beat amplitude, and excitation-contraction delay. Samples were omitted if the electrical signal had an insufficient signal-to-noise ratio or if hiPSC-CM were not pacing at the desired frequency. To account for slight differences between individual plates or hiPSC-CM lot variability, vehicle controls (DMSO) were included and recorded on each MEA plate.

**Figure 1.**
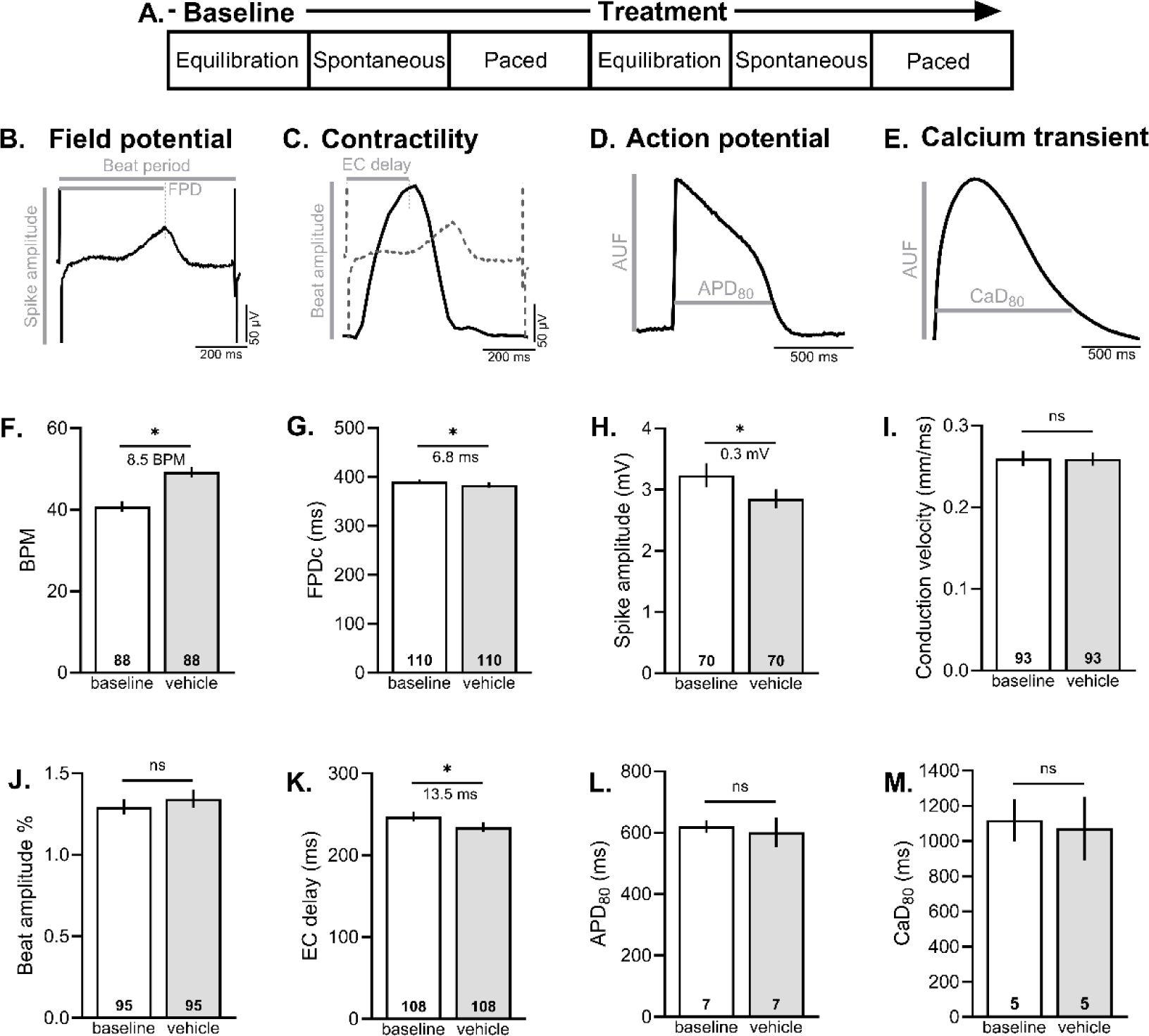
Experimental protocol. **A)** hiPSC-CM were equilibrated (15-20 min), and then basal cardiac metrics were measured during spontaneous beating or in response to external pacing. Thereafter, hiPSC-CM were treated with a specified chemical for 15-20 min, and then cardiac metrics were measured again. Representative traces and fiducial markers are shown for **B)** field potential recording, **C)** contractility, **D)** optical action potential and **E)** intracellular calcium transient. **F-K)** MEA parameters were compared between baseline and 0.1% DMSO (vehicle) to establish threshold values for chemical treatment studies. **L-M)** Action potentials and calcium transients were also compared between baseline and 0.1% DMSO. *Number of independent replicates reported in each bar. Mean and 95% confidence interval are reported; T-test with *p<0.05. BPM recorded during spontaneous rhythm, all other parameters reported in response to pacing. APD_80_: action potential duration at 80% repolarization; AUF: arbitrary unit of fluorescence; BPM: beats per minute; CaD_80_: calcium transient duration at 80% recovery, EC delay: excitation-contraction delay time, FPDc: field potential duration, rate corrected*.

### Fluorescent imaging

hiPSC-CM were loaded with a calcium indicator dye (10 μM Fluo 4-AM, Thermo Fisher Scientific) or a potentiometric dye (1:1000 FluoVolt, Thermo Fisher Scientific) for 20 min at room temperature (22°C), then washed in dye-free Tyrode salt solution (Sigma-Aldrich), as previously described (Cooper et al. 2021). Cell monolayers were treated for 15-20 min at room temperature. Pace-induced calcium transients and action potential recordings were acquired using a Nikon TiE microscope system, equipped with 470 nm excitation LED (SpectraX, Lumencor), 505–530 nm emission filter, and Photometrix 95B sCMOS back-illuminated camera. Cardiac monolayers were paced using field stimulation (0.2–1 Hz). Optical signals were analyzed using custom software (Cooper et al. 2021; Jaimes et al. 2016). Cardiac parameters included: calcium transient activation time, duration time (CaD_30_ or CaD_80_; activation to 30% or 80% recovery), Tau (decay time constant), and action potential duration time (APD_30_ or APD_80_; activation to 30% or 80% repolarization time).

### Statistical analysis

Results were reported as the mean and 95% confidence interval, unless otherwise indicated. Statistical analysis was performed using either a Student’s t-test (two groups) or one-way analysis of variance (ANOVA) with Holm-Sidak multiple comparison test (three or more groups), as indicated in each figure legend. An adjusted p-value <0.05 was considered statistically significant. Replicates are defined in each figure.

## RESULTS

### Experimental protocol

We first measured variations in hiPSC-CM physiology under baseline conditions and following exposure to 0.1% DMSO – which was used as a vehicle in subsequent bisphenol studies (**Figure 1F-M**). Application of DMSO had no effect on cardiomyocyte conduction velocity (*baseline:* 0.26 [0.25,0.27], *vehicle:* 0.259 [0.25,0.27] mm/ms) or beat amplitude (*baseline:* 1.29 [1.25,1.34], *vehicle:* 1.34 [1.29,1.40] %). Further, the application of DMSO did not have an effect on action potential duration (APD_80_ *baseline:* 620 [599,641], *vehicle:* 602 [554,650] ms) or calcium transient duration (CaD_80_ *baseline:* 1119 [999,1238], *vehicle:* 1071 [891,1251] ms), as assessed by fluorescent imaging. Supplementing cell culture media with 0.1% DMSO slightly shifted the spontaneous beating rate (*baseline:* 40.8 [39.5,42.1], *vehicle:* 49.3 [48.0,50.6] BPM, p<0.0001), FPDc (*baseline:* 390.4 [385.1,395.6], *vehicle:* 383.6 [378.3,388.8] ms, p<0.001), depolarization spike amplitude (*baseline:* 3.24 [3.04,3.43], *vehicle:* 2.85 [2.70,3.00] mV, p<0.0001), and excitation-contraction delay (*baseline:* 247.0 [241.2,252.9], *vehicle:* 234.5 [228.8,240.1] ms, p<0.001). To take these variations into account, cardiac measurements were compared between DMSO and bisphenol chemical exposure.

### Acute effects of E2 and BPA on cardiac electrophysiology

For toxicity studies, we first identified the timeframe in which BPA or E2 alter cardiac electrophysiology (**Figure 2**). Previous studies report rapid effects of BPA, with no discernable effect on cardiomyocyte viability up to 100 μM (Hyun et al. 2021; Posnack et al. 2014; Ramadan et al. 2018). hiPSC-CM spontaneous beating rate remained relatively stable after exposure to low doses of BPA (1-100 nM), while 30 μM BPA transiently increased the beating rate (*20 minutes vehicle:* 62.0 [57.1,66.9], *BPA*: 77.1 [72.9,81.4] BPM, p<0.0001) with effects waning over time (**Figure 2A**). Similarly, 30 μM E2 rapidly increased the beating rate within 5 minutes, but thereafter all spontaneous activity ceased until approximately 24 hours later (**Figure 2B**). We also noted an immediate effect of BPA and E2 on the rate-corrected field potential duration (FPDc), which had a nonlinear dose-response. At lower doses, FPDc was slightly longer following acute BPA (*20 minutes vehicle:* 328.3 [319.9,336.7], *100 nM*: 342.0 [332.6,351.4] ms, p<0.05) or E2 treatment (*20 minutes 1 nM:* 341.2 [333.9,348.5], *100 nM*: 338.9 [331.8,345.9] ms, p<0.05; **Figure 2C-D**). While at higher doses, FPDc shortened with acute BPA (*20 minutes 30 μM:* 300.3 [294.3,306.3] ms, p<0.0001) or E2 treatment (*30 μM:* 273.4 [266.3,280.6] ms, p<0.0001). Notably, these effects were reversible after the removal of BPA-or E2-supplemented media (data not shown). Others have reported arrhythmias (triggered activity) in primary cardiomyocytes after BPA exposure (Belcher et al. 2011; Yan et al. 2011), but we did not observe rhythm or propagation irregularities in BPA-exposed hiPSC-CM (**Figure 2E**). However, acute 30 μM E2 treatment significantly reduced the propagation consistency (*20 minutes vehicle:* 94.3 [89.3,99.3], *30 μM:* 52.1 [38.4,65.8] %, p<0.0001)

**Figure 2.**
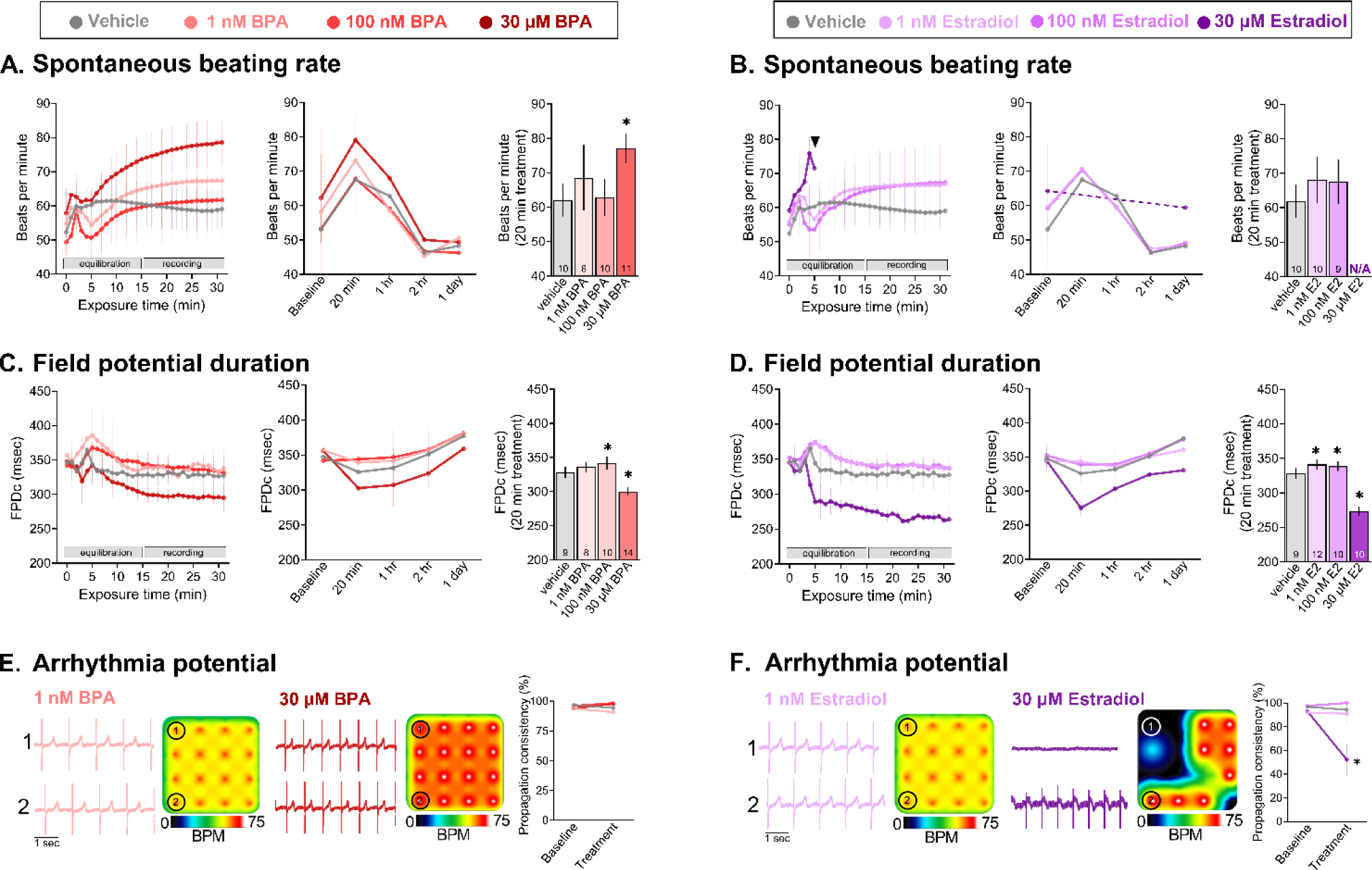
Time-dependent effects of BPA and E2. **A-B)** Spontaneous beating rate following BPA or E2 exposure. ANOVA with multiple comparisons test, *p<0.05 compared with vehicle (0.1% DMSO). **C-D)** Field potential duration (rate-corrected with Fridericia formula) following BPA or E2 exposure. ANOVA with Holm-Sidak multiple comparisons test, *p<0.05 compared with vehicle (0.1% DMSO). **E-F)** Arrhythmia potential as determined by propagation consistency across the microelectrode array. Raw signals from two individual electrodes are shown, as well as a heatmap of the spontaneous beating rate across the 16-electrode array. *Paired t-test, *p<0.05 compared with baseline recording on same electrode. Mean and 95% confidence interval are reported. BPM recorded during spontaneous beating, FPDc recorded in response to 1.5 Hz pacing frequency. N/A: Not assessed due to loss of spontaneous beating activity*.

### Dose-dependent effects of E2 and bisphenol chemicals on hiPSC-CM electrophysiology

After confirming that the cardiac effects of E2 and BPA occur within an acute timeframe (∼20 minutes), we next sought to compare the potency of bisphenol chemicals. The spontaneous beating rate was inconsistent between treatment groups, with loss of activity observed upon exposure to higher concentrations of E2, BPA, and BPF – but not BPS (**Figure 3**). As such, cardiac parameters were assessed in response to external pacing to account for rate-dependent adaptations.

**Figure 3.**
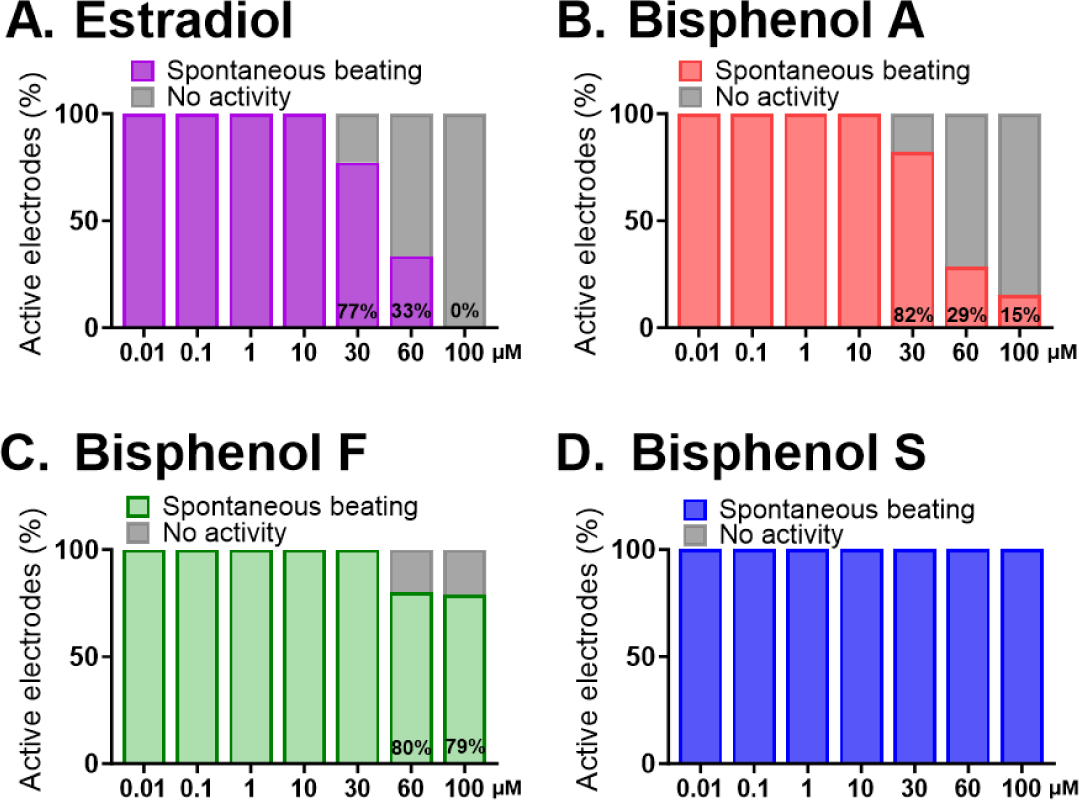
Effect of E2 and bisphenol chemicals on hiPSC-CM spontaneous beating. hiPSC-CM spontaneous beating activity slowed after exposure to higher concentrations of **A)** E2, **B)** BPA, **C)** BPF, but not BPS **(D)**. Values reported as the percentage of cardiomyocytes with spontaneous activity, as recorded from cell monolayers plated atop microelectrode arrays. n=11-40 individual cardiomyocyte preparations per dose, per treatment.

Since BPA inhibits voltage-gated Na^+^ channels through interaction with the anesthetic receptor site (O’Reilly et al. 2012; Prudencio et al. 2021), we examined the direct effect of E2 and bisphenol chemicals on the depolarization spike amplitude which corresponds to Na^+^ influx (**Figure 4**). Spike amplitude remained unchanged at lower concentrations of E2 and BPA, but decreased at higher micromolar concentrations (≥10 μM; **Figure 4A-C**). Compared with vehicle control, E2 and BPA reduced the spike amplitude by 14-17.8% at 30 μM (*vehicle*: 2.88 [2.73,3.02], *E2*: 2.35 [1.95,2.74], *BPA:* 2.45 [2.23,2.68] mV) and by 85.3-87.1% at 60 μM (*E2:* 0.42 [0.33,0.51], *BPA:* 0.37 [0.33,0.41] mV; **Figure 4B-C**). This aligns with the reported half-maximal inhibitory concentration (23-56 μM) for BPA on sodium channel current (Hyun et al. 2021; O’Reilly et al. 2012; Prudencio et al. 2021). Comparatively, BPF and BPS exposure had a less pronounced effect on the depolarization spike amplitude. Next, we measured conduction velocity across the microelectrode array – as a reduced spike amplitude is expected to delay electrical propagation within a cardiomyocyte monolayer. Conduction velocity slowed after acute exposure to E2 or BPA, with less pronounced effects observed with BPF or BPS exposure (**Figure 4D-E**). To account for any slight differences between hiPSC-CM experiments due to lot variability or plating density, both raw (**Figure 4B, D**) and percent change data are reported (**Figure 4C, E**) – wherein the latter accounts for differences in baseline measurements before applying a chemical treatment.

**Figure 4.**
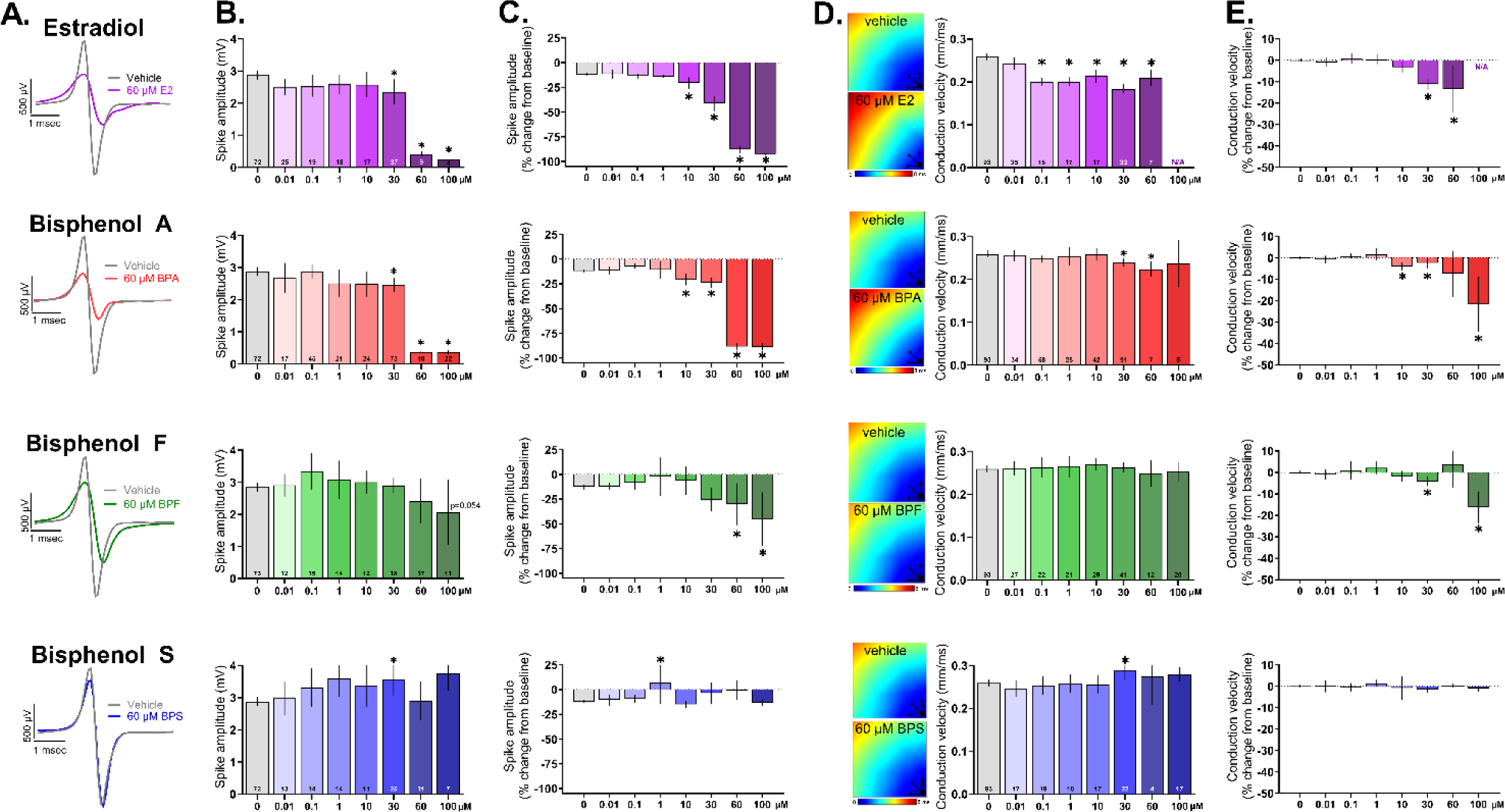
Effect of E2 and bisphenol chemicals on depolarizing spike amplitude and conduction velocity. **A)** Representative traces of depolarizing spike recorded from hiPSC-CM, paced at 1.5 Hz, following acute (15 min) exposure to 60 μM E2, BPA, BPF, or BPS. Note: the y-axis is truncated in order to view the small depolarizing spike resulting from E2 or BPA exposure. **B)** Spike amplitude, chemical dose response. **C)** Percent change in spike amplitude after chemical exposure, relative to matched baseline recording. **D)** <u>Left:</u> Activation maps recorded from a monolayer of hiPSC-CM, paced at 1.5 Hz, following acute (15 min) exposure to 60 μM E2, BPA, BPF, or BPS. Activation is initiated at the pacing electrode in lower right corner, as indicated. <u>Right:</u> Conduction velocity, chemical dose response. **E)** Percent change in conduction velocity after chemical exposure, relative to matched baseline recording. *Number of independent replicates reported in each bar. Mean and 95% confidence interval are reported; ANOVA with multiple comparisons test, *p<0.05 relative to vehicle control. N/A: Not assessed due to loss of activity. Conduction velocity was only measured on hiPSC-CM with consistent electrical propagation*.

Previous studies demonstrated that BPA shortens the extracellular FPD (analogous to the electrocardiogram QT interval), which may be related to its inhibitory effect on voltage-gated Ca^2+^ channels (Hyun et al. 2021; Prudencio et al. 2021). At a pacing frequency of 1.5 Hz, the rate-corrected FPD shortened upon exposure to higher concentrations of E2, BPA, and BPF (**Figure 5**). As an example, compared with vehicle control, the FPDc shortened by 11.8-18 % at 30 μM dose (*vehicle:* 383.6 [378.3, 388.8], *E2*: 314.3 [307.5,321.1], *BPA*: 333.7 [329.2, 338.2], *BPF:* 338.3 [329.8,346.8] msec; **Figure 5A-C**). In comparison, BPS exposure had no measurable effect on FPDc at any of the tested concentrations (0.01-100 μM). Both the raw (**Figure 5B, D**) and percent change FPDc data are reported (**Figure 5C**) to account for any slight differences in hiPSC-CM phenotype between experiments. In a subset of studies, we also tested the electrical restitution properties of hiPSC-CM by incrementally increasing the pacing frequency to assess rate-dependent adaptations. In control preparations, the FPDc shortened by 14.3-16.2% as the pacing frequency increased from 1 to 2 Hz, with a clearly defined negative slope (**Figure 5D**). This rate-dependent adaptation was maintained in the presence of BPS (0.1-60 μM), but exposure to elevated concentrations of E2, BPA, or BPF resulted in a flattened electrical restitution curve. As an example, vehicle control samples had a restitution slope of −57 to −71 across all replicates, which flattened significantly compared to 30 μM E2 (−1 slope), BPA (−31 slope), and BPF (−51 slope; **Figure 5D**). FPDc measurements were inconsistent at 100 μM E2 and BPA, as hiPSC-CM became unresponsive to electrical stimulation.

**Figure 5.**
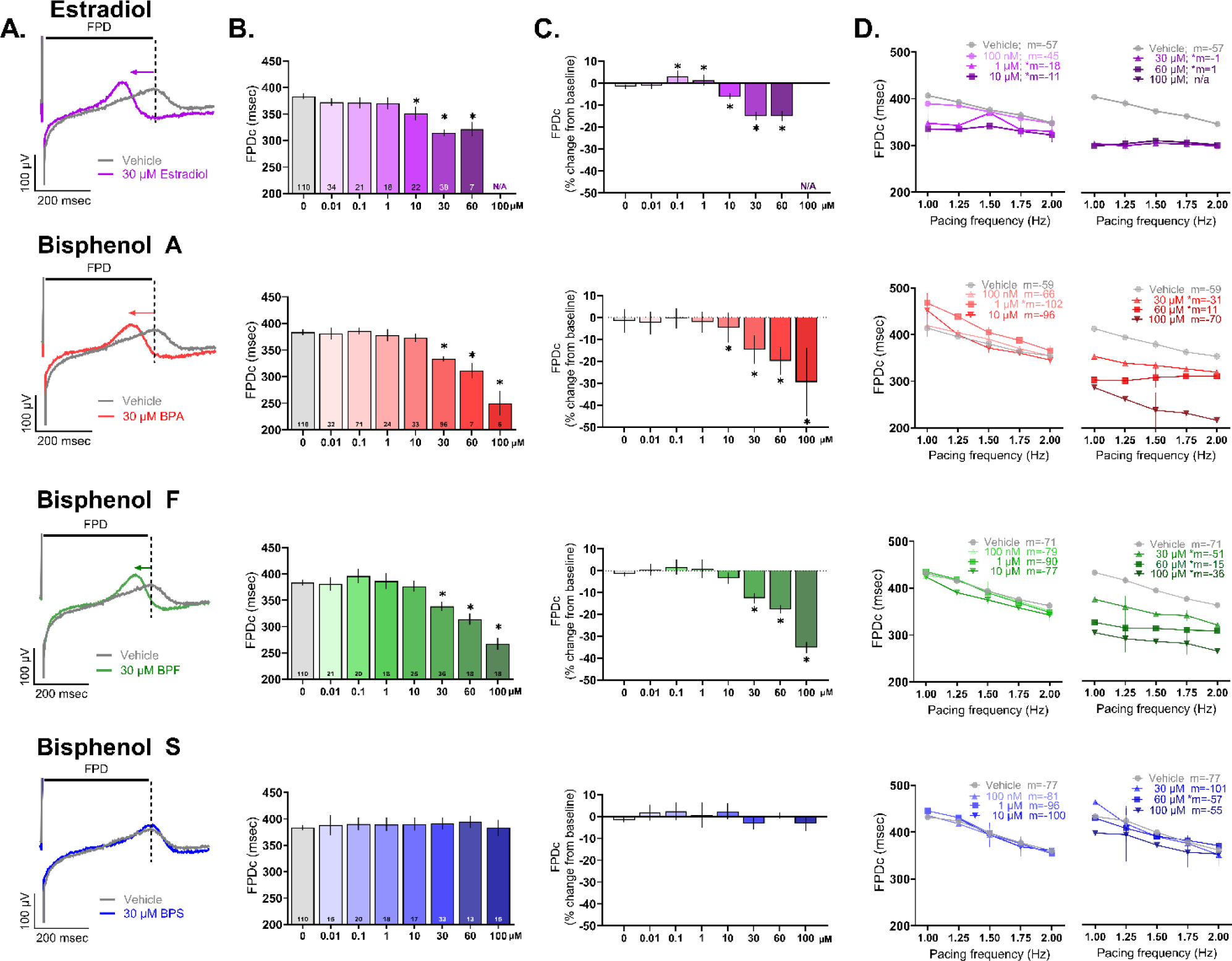
Effect of E2 and bisphenol chemicals on extracellular field potential duration (FPD). **A)** Representative extracellular field potential traces recorded from hiPSC-CM, paced at 1.5 Hz, following acute (15 min) exposure to 30 μM E2, BPA, BPF, or BPS. Note: the y-axis is truncated in order to zoom in on the T-wave (dotted line). **B)** FPDc, chemical dose response. **C)** Percent change in FPDc after chemical exposure, relative to matched baseline recording. **D)** Electrical restitution with pacing rate increased stepwise from 1-2 Hz; slope (“m”) is indicated for each chemical. *Number of independent replicates reported in each bar. Mean and 95% confidence interval are reported; ANOVA with multiple comparisons test, *p<0.05 relative to vehicle control. N/A: Not assessed due to loss of activity. Samples omitted if not responsive to the desired pacing frequency*

In a subset of studies, hiPSC-CM were electrically induced to increase cell-to-electrode coupling to record local extracellular action potentials (Hayes et al. 2019). The dose response patterns were similar to FPD results, wherein the action potential duration (APD) shortened following exposure to increasing concentrations of E2, BPA, and BPF. As an example, compared to vehicle control, APD_90_ shortened by 7.9-20.2% at 30 μM dose (*vehicle:* 396.7 [388.1, 405.3], *E2*: 316.7 [309.8,323.6], *BPA*: 365.3 [352.6,325.3], *BPF:* 360.3 [345.4,375.2] msec; 1.5 Hz pacing frequency; **Figure 6A, B**). Comparatively, BPS had no effect on APD_90_ at any of the tested concentrations (0.01-100 μM). We also tested rate-dependent adaptations, wherein APD_90_ shortened by 11.4% (−101 slope) in control cell preparations as the pacing frequency increased from 1 to 2 Hz (**Figure 6C**). Conversely, higher concentrations of E2, BPA, and BPF produced a flattened electrical restitution curve compared with the control – while exposure to lower doses of BPS (0.01-10 μM) resulted in a slightly steeper curve. We also analyzed optical action potentials using a voltage-sensitive dye, which yielded similar results (**Figure 6D**, 0.5 Hz frequency). APD_80_ shortening was observed after exposure to E2 (*baseline:* 867 [791,943], *30 μM E2:* 566 [484,649] msec), BPA (*baseline:* 783 [680,886], *30 μM BPA:* 568 [510,626] msec), and BPF (*baseline:* 862 [763,960], *30 μM BPF*: 688 [637,738] msec) – but not BPS (*baseline:* 757 [639,876], *30 μM BPS:* 742 [627,856] msec).

**Figure 6.**
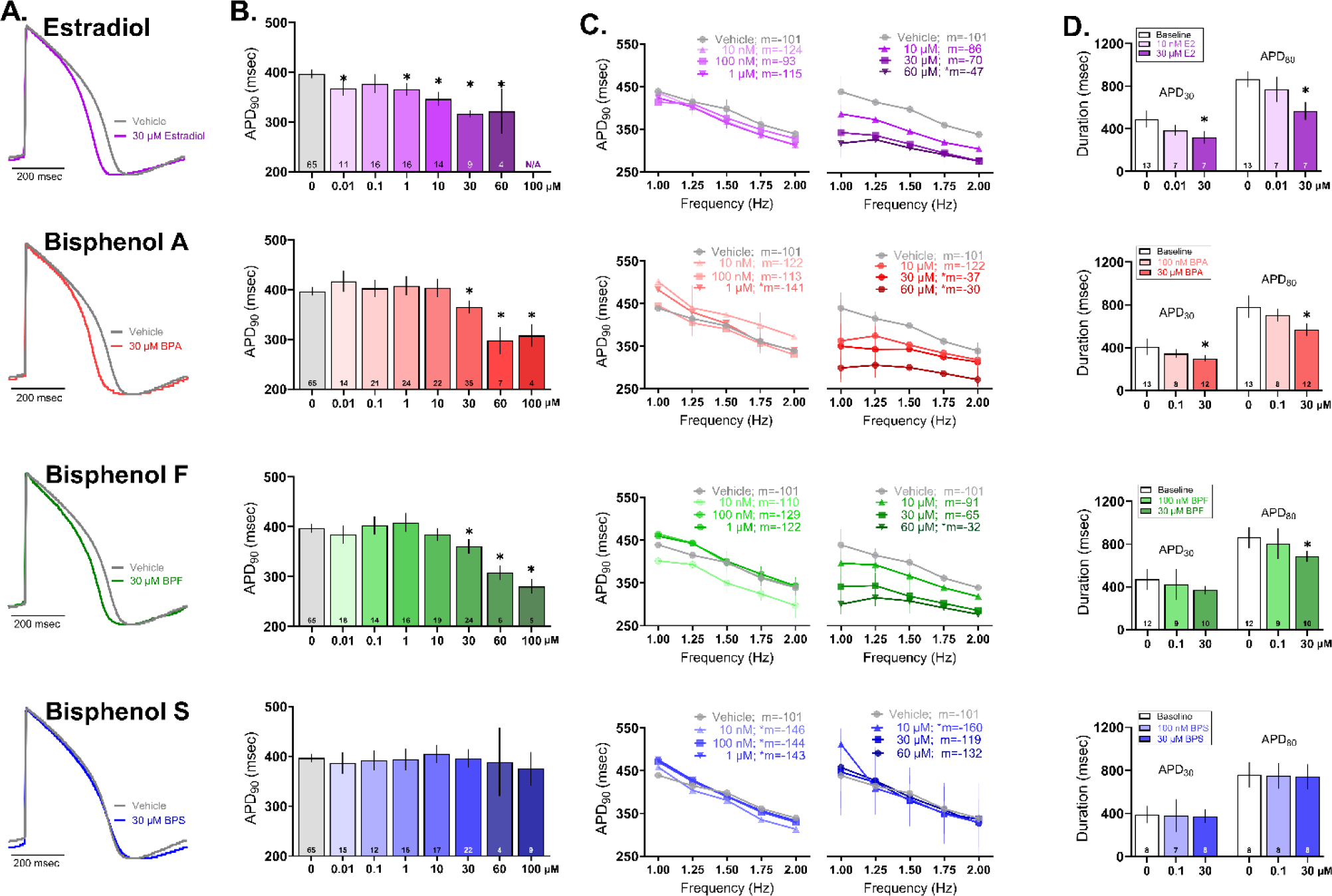
Effect of E2 and bisphenol chemicals on cardiac action potential duration. **A)** Representative action potential traces recorded from hiPSC-CM following 15 min exposure to 30 μM E2, BPA, BPF, or BPS (1.5 Hz pacing frequency, 37°C). **B)** Local extracellular action potential duration at 90% repolarization (APD_90_), dose response. **C)** Electrical restitution with pacing rate increased stepwise from 1-2 Hz; slope “m” is indicated for each chemical. **D)** Action potential duration at 30% and 80% repolarization, measured optically using a voltage-sensitive dye (0.5 Hz pacing frequency, room temperature). *Number of independent replicates reported in each bar. Mean and 95% confidence interval are reported; ANOVA with multiple comparisons test, *p<0.05 relative to vehicle control (A-C) or baseline measurement preceding chemical exposure (D). N/A: Not assessed due to loss of activity. Samples omitted if not responsive to the desired pacing frequency or if optical signals were of insufficient quality for analysis*.

### Dose-dependent effects of E2 and bisphenol chemicals on calcium handling and contractility

Prior studies suggest that E2 and BPA interfere with intracellular calcium transients, which may be a consequence of L-type calcium channel inhibition and/or the phosphorylation of key regulatory proteins (Cooper and Posnack 2022; Gao et al. 2013; Hyun et al. 2021; Prudencio et al. 2021). To evaluate such effects, we recorded calcium transients from hiPSC-CM using an indicator dye (Fluo4; **Figure 7A**). All treated samples showed rate-dependent adaptations, wherein the calcium transient duration (CaD) shortened as the pacing frequency was increased (0.2-1 Hz) – but to varying degrees. Rate-dependent shortening of CaD_80_ was slightly blunted with a flattened curve observed after exposure to E2 (*baseline:* 48.9%, *30 μM E2:* 42.7% shortening) – but a greater change upon exposure to BPA (*baseline:* 46.1%, *30 μM BPA:* 35.5% shortening), BPF (*baseline:* 42.5%, *30 μM BPF:* 33% shortening), or BPS (*baseline:* 48.4%, *30 μM BPS:* 37.2% shortening; **Figure 7B**).

**Figure 7.**
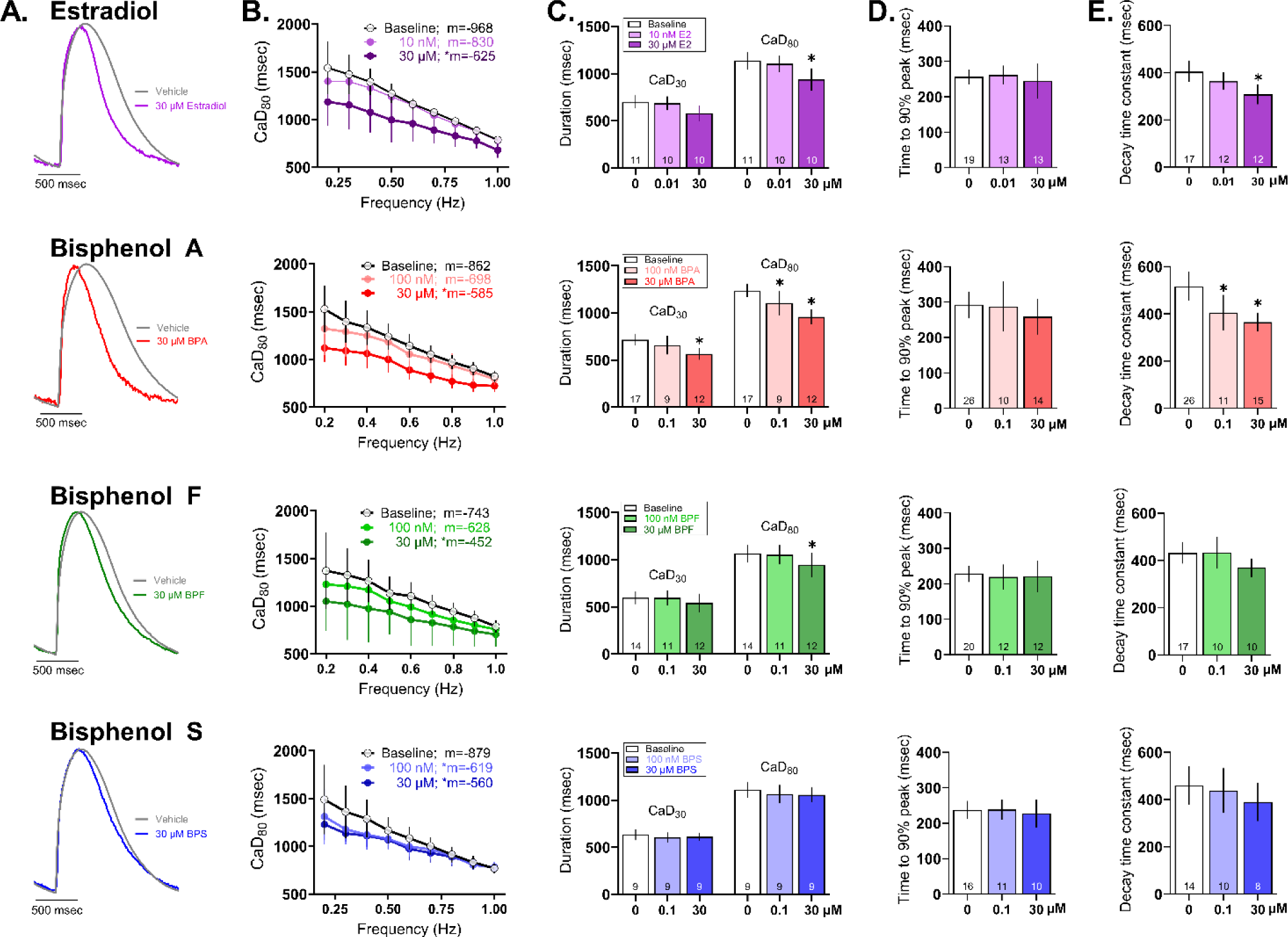
Effect of E2 and bisphenol chemicals on intracellular calcium transients. **A)** Representative calcium transients recorded from hiPSC-CM following 15 min exposure to 30 μM E2, BPA, BPF, or BPS. Calcium transients were measured optically using a calcium-indicator dye (0.5 Hz pacing frequency, room temperature). **B)** Calcium transient duration at 80% (CaD_80_) as the pacing rate was increased stepwise from 0.2-1 Hz; slope “m” is indicated for each chemical. **C)** CaD_30_ and CaD_80_, **D)** Time to 90% peak fluorescence, **E)** Decay time constant all measured at 0.5 Hz frequency. *Number of independent replicates reported in each bar. Mean and 95% confidence interval are reported; ANOVA with multiple comparisons test, *p<0.05 relative to baseline measurements preceding chemical treatment. Samples omitted if not responsive to the desired pacing frequency or if optical signals were of insufficient quality for analysis*.

At an intermediate pacing frequency (0.5 Hz), the calcium transient duration at 80% recovery (CaD_80_) was shorter after exposure to 30 μM E2 (*baseline:* 1137 [1042,1231], *E2:* 938.6 [821.6,1056] msec), BPA (*baseline:* 1237 [1167,1306], *BPA*: 958.3 [881.8,1035] msec), and BPF (*baseline:* 1068 [974.4,1161], *BPF:* 945.6 [815.4,1078] msec) – but BPS had no measurable effect on CaD_80_ (**Figure 7C**). The timing of calcium-induced calcium release did not appear to be altered, as assessed by the time to reach 90% of the calcium transient peak (measured via fluorescence, **Figure 7D**). Instead, the effects on calcium handling were related to the recovery phase (assessed by the decay time constant, **Figure 7E**) reflecting the time required for calcium sequestration into the sarcoplasmic reticulum or extrusion from the cell. CaD_80_ shortening may be caused by increased phospholamban phosphorylation which enhances sarcoplasmic reticulum calcium-ATPase activity (Gao et al. 2013) and/or less inward calcium current which diminishes calcium release from the sarcoplasmic reticulum and shortens the calcium transient amplitude (Prudencio et al. 2021; Ramadan et al. 2018). Such differences in calcium handling were rate-dependent and masked at faster pacing frequencies.

Next, we assessed the impact of chemical exposure on the magnitude and timing of cardiomyocyte contraction. We anticipated that perturbations in calcium handling would reduce contractility in BPA-treated hiPSC-CM, as previously reported in rodent heart and atrial preparations (Pant et al. 2011; Posnack et al. 2015). Indeed, the contractile beat amplitude decreased following exposure to increasing concentrations of E2 (*baseline:* 1.32% [1.29,1.34], *60 μM:* 0.56% [0.54,0.57], *100 μM:* 0.53% [0.50,0.55]), BPA (3*0 μM:* 1.15% [1.07,1.23], *60 μM:* 0.65% [0.61,0.69], *100 μM:* 0.57% [0.52,0.61]), or BPF (*100 μM:* 0.97% [0.71,1.22]) – while BPS had no effect on contractility (**Figure 8A-C**). We also measured the coupling time between the electrical and contractile signal and noted a significant 46-61% longer coupling time after exposure to 60-100 μM E2 or BPA (**Figure 8D, E**). Excitation-contraction coupling time was unaffected by either BPF or BPS.

**Figure 8.**
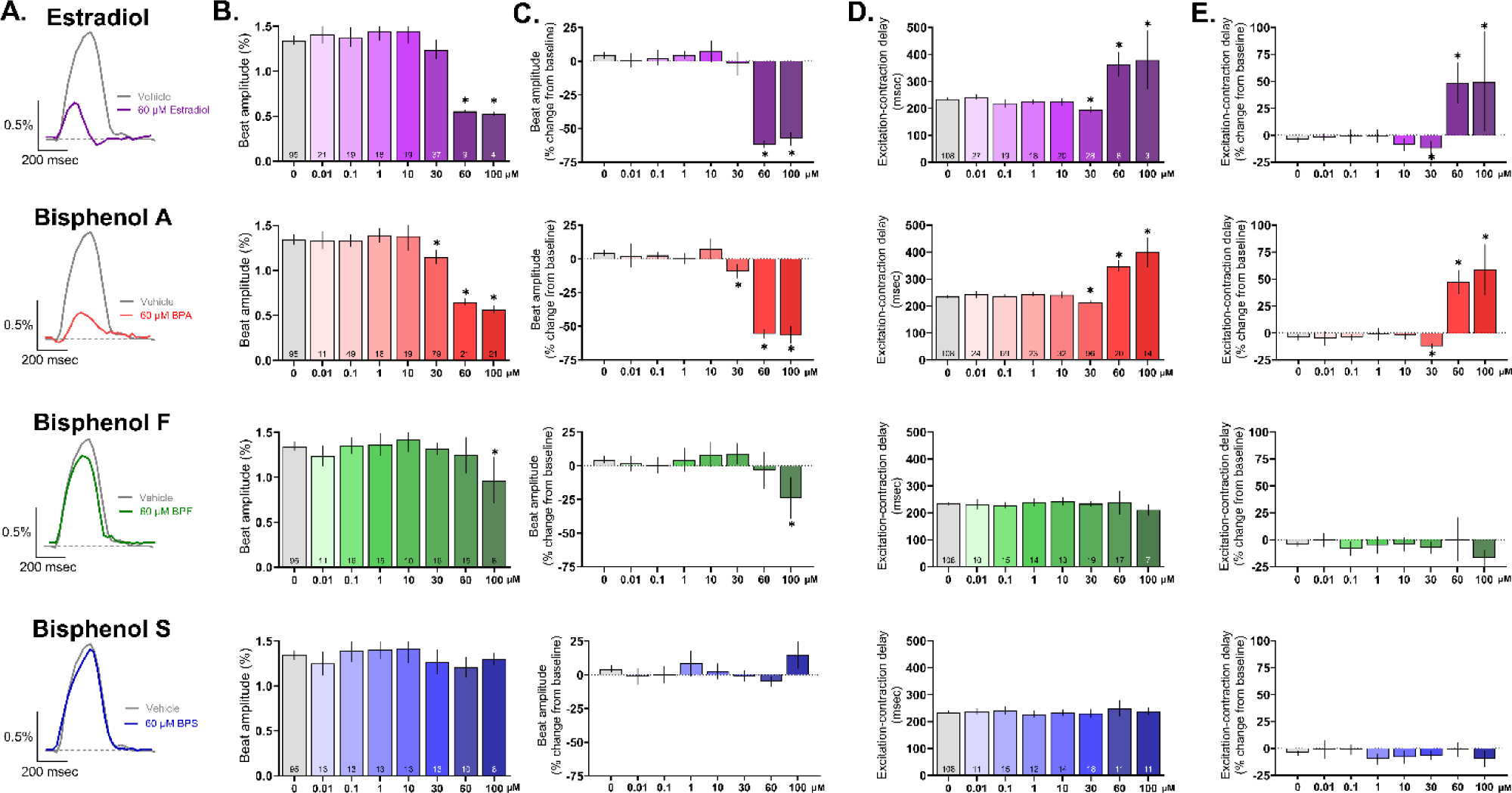
Effect of E2 and bisphenol chemicals on cardiomyocyte contraction. **A)** Representative recordings of hiPSC contraction and relaxation (impedance-based measurement) following 15 min exposure to 60 μM E2, BPA, BPF, or BPS. **B)** Beat amplitude, dose response. **C)** Percent change in beat amplitude after chemical exposure, relative to matched baseline recording. **D)** Excitation-contraction coupling time, dose response. **E)** Percent change in excitation-contraction coupling time, relative to matched baseline recording. *Number of independent replicates reported in each bar. Mean and 95% confidence interval are reported; ANOVA with multiple comparisons test, *p<0.05 relative to vehicle control (0.1% DMSO). Samples omitted if not responsive to the desired pacing frequency or if the contraction amplitude was too small to measure accurately*.

### BPA-induced effects are exaggerated by co-administration with E2 or calcium channel antagonist

Several studies have suggested that BPA cardiotoxicity is related to its interaction with estrogen receptors and/or the inhibition of voltage-gated calcium channels (Belcher et al. 2011; Deutschmann et al. 2013; Feiteiro et al. 2018; Michaela et al. 2014; Prudencio et al. 2021; Yan et al. 2011). Accordingly, we utilized pharmacological tools to assess the cellular targets of E2 and BPA, and to determine whether these agents exert an additive effect. A single concentration of E2 and BPA was used (30 μM), which was efficacious in our prior results and aligns with the half-maximal inhibitory concentration of BPA on voltage-gated calcium channels (Hyun et al. 2021; Prudencio et al. 2021). In this subset of experiments, 30 μM BPA and 30 μM E2 increased the spontaneous beating rate to a similar degree (+25.6% and +35.7%, respectively; **Figure 9**). Next, we evaluated whether this effect could be attenuated by pretreating hiPSC-CM with an L-type calcium channel (LTCC) agonist (Bay K8644), which effectively reduced the effects of BPA and E2 on beating rate (**Figure 9A,B**). Pretreatment with an LTCC antagonist (verapamil) exaggerated the effect of BPA on beating rate, but not E2 – suggesting a different mechanism of action.

**Figure 9.**
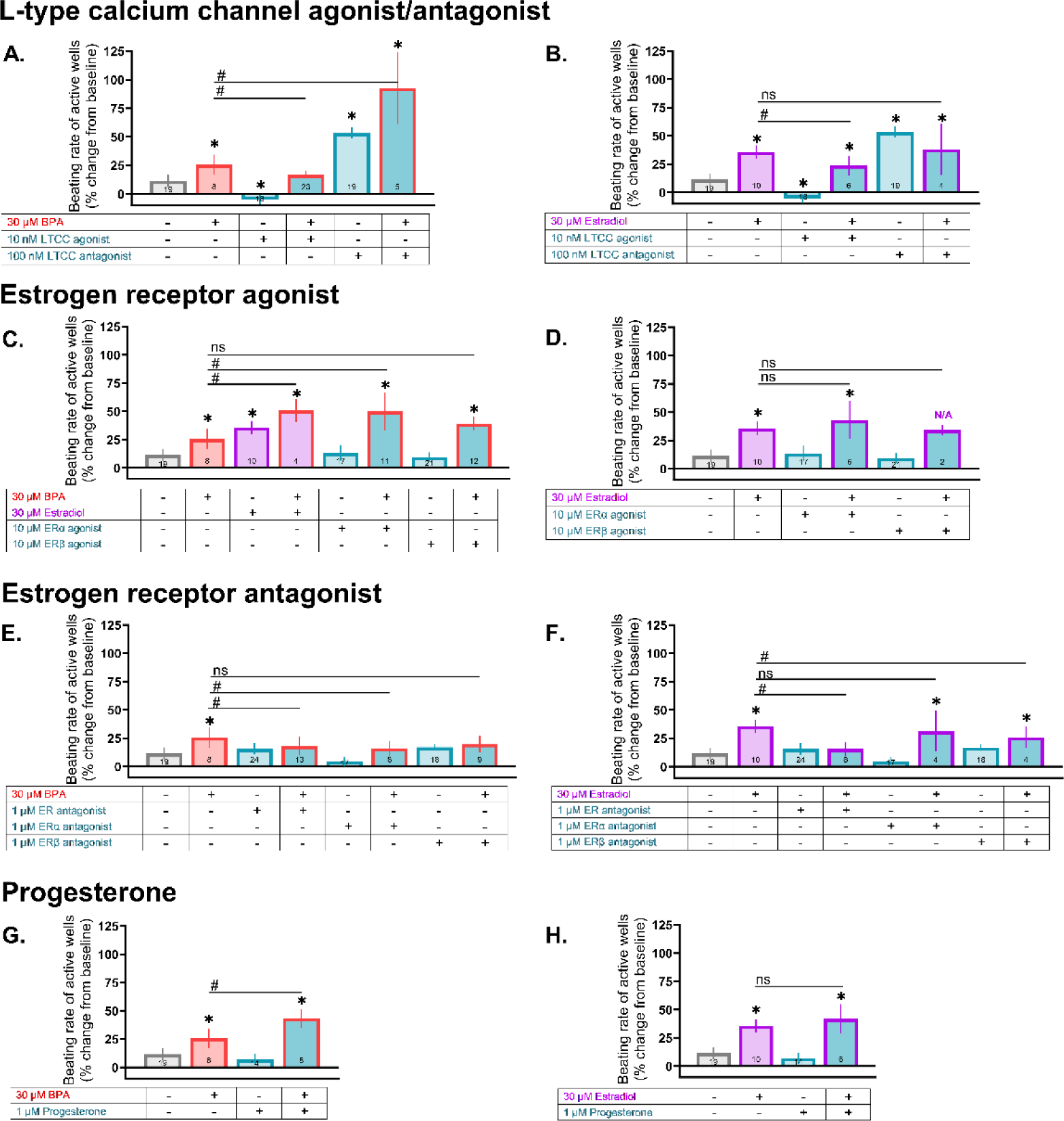
BPA-induced changes on beating rate are exaggerated by co-administration with E2 or calcium channel antagonist. hiPSC-CM were pretreated with a pharmacological agent (15 min) followed by exposure to either 30 μM BPA or 30 μM E2. **A,B)** Pretreatment with an LTCC agonist (Bay K8644) or antagonist (verapamil). **C,D)** Pretreatment with at ERα agonist (PPT) or ERβ agonist (DPN). **E,F)** Pretreatment with a non-specific ER antagonist (ICI 182,780), ERα antagonist (MPP), or ERβ antagonist (PHTPP). **G,H)** Pretreatment with progesterone. *Number of independent replicates reported in each bar. Values reported as a percent change from baseline measurement, immediately preceding chemical treatment (mean and 95% confidence interval). ANOVA with multiple comparisons test, *p<0.05 relative to vehicle control (0.1% DMSO), #p<0.05 relative to either 30 μM BPA alone or 30 μM E2 alone. Samples omitted if spontaneous beating activity ceased. N/A: not applicable due to loss of spontaneous activity across multiple samples, which prevented statistical analysis*.

Studies also suggest that BPA cardiotoxicity is amplified by E2 – likely through estrogen receptor signaling (Belcher et al. 2011; Yan et al. 2011, 2013), and curtailed by progesterone (Ma et al. 2017). In our hiPSC-CM model, BPA+E2 caused an exaggerated effect on the spontaneous beating rate (+50.8% above baseline), as did BPA+ERα agonist (PPT; +50% above baseline) – but not BPA+ERβ agonist (DPN; **Figure 9C**). Pretreatment with either a non-specific ER antagonist (ICI 182,780) or ERα antagonist (MPP) slightly reduced the beating rate effects of BPA – but pretreatment with an ERβ antagonist did not (PHTPP; **Figure 9E**). Unexpectedly, progesterone pretreatment exaggerated the effects of BPA on beating rate (+43% above baseline; **Figure 9G**). A few notable differences were observed when compared to E2. First, co-administration with either an ERα or ERβ agonist did not cause an additive effect on beating rate (**Figure 9D**). Notably, when E2 was added with an ERβ agonist, loss of spontaneous electrical activity was observed for most replicates. Second, the effects of E2 were attenuated by an ERβ antagonist, but not an ERα antagonist (**Figure 9F**). Third, the effects of E2 were not altered by progesterone pretreatment (**Figure 9H**).

Finally, we assessed whether these same pharmacological tools prevented FPDc shortening in BPA or E2 treated hiPSC-CM. In this subset of experiments, 30 μM BPA and 30 μM E2 shortened the FPDc to a similar degree (−13.4% and −17%, respectively), while BPA+E2 exaggerated this effect (−21.4%; **Figure 10E, F**). Pretreatment with an LTCC agonist (Bay K8644) reduced the effects of both BPA and E2 on FPDc, and an LTCC antagonist (verapamil) exaggerated this effect when co-administered with BPA, but not E2 (**Figure 10A-D**). FPDc shortening was (slightly) exaggerated when BPA was combined with an ERα agonist (PPT; −16.4%), non-specific ER antagonist (ICI 182,780; −22.3%), or progesterone (−16.7%) - and reduced when combined with an ERβ agonist (DPN; −12.7%) or ERα antagonist (MPP; −7.7%; **Figure 10**). E2-treated cells yielded slightly different results; first, an exaggerated shortening of FPDc was not observed with either the ERα agonist (PPT) or progesterone (**Figure 10H, P**). Second, the non-specific ER antagonist reduced the effects of E2 (**Figure 10L**). These findings suggest that E2 and BPA may act on different targets, or may differ in potency for L-type calcium channels and/or estrogen receptors.

**Figure 10.**
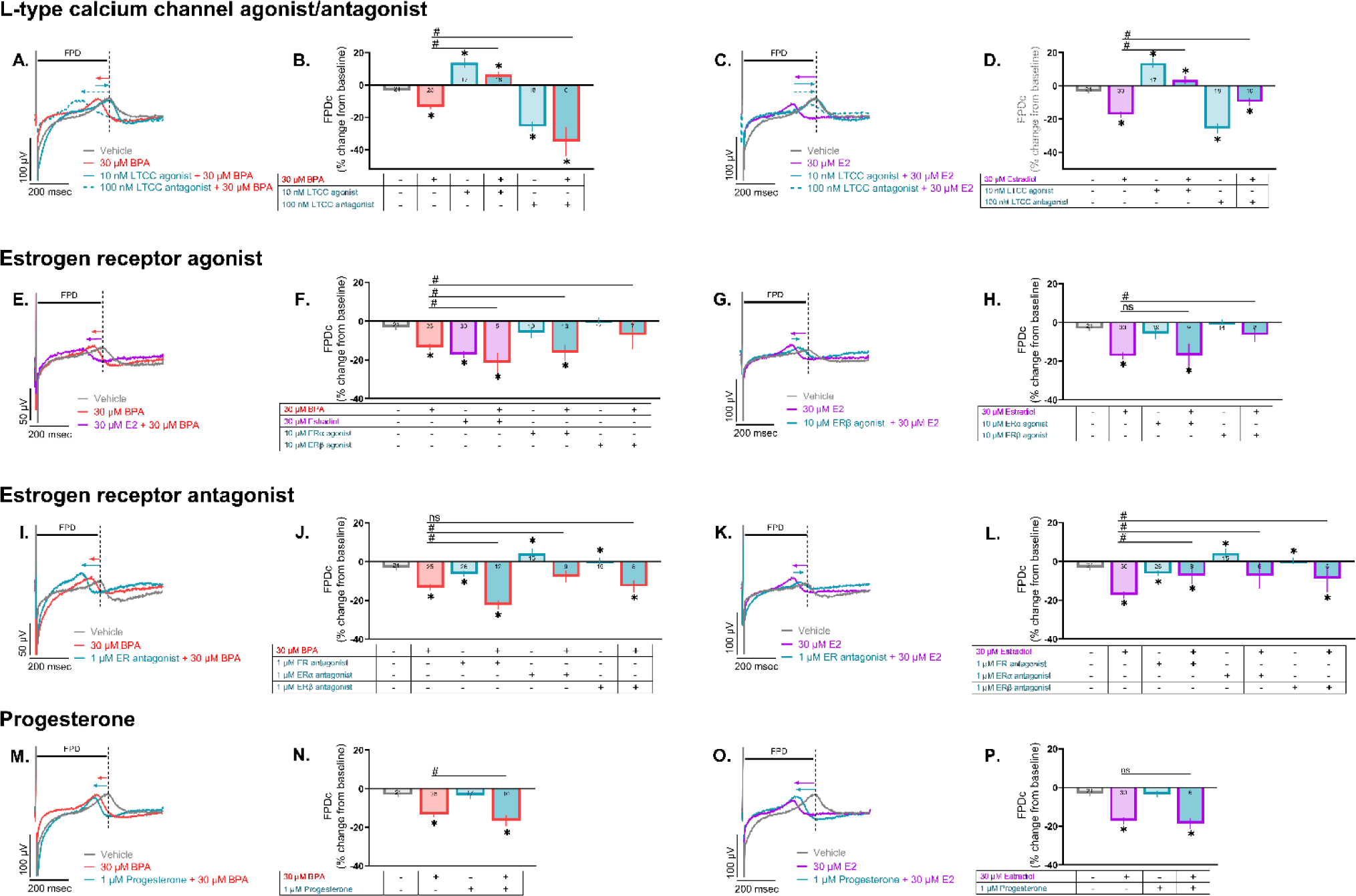
BPA-induced changes on FPDc are exaggerated by co-administration with E2 or calcium channel antagonist. hiPSC-CM were pretreated with a pharmacological agent (15 min) followed by exposure to either 30 μM BPA or 30 μM E2. Field potentials recorded in response to 1.5 Hz pacing. **A-D)** Pretreatment with an LTCC agonist (Bay K8644) or antagonist (verapamil). **E-H)** Pretreatment with at ERα agonist (PPT) or ERβ agonist (DPN). **I-L)** Pretreatment with a non-specific ER antagonist (ICI 182,780), ERα antagonist (MPP), or ERβ antagonist (PHTPP). **M-P)** Pretreatment with progesterone. *Number of independent replicates reported in each bar. Values reported as a percent change from baseline measurement, immediately preceding chemical treatment (mean and 95% confidence interval). ANOVA with multiple comparisons test, *p<0.05 relative to vehicle control (0.1% DMSO), #p<0.05 relative to either 30 μM BPA alone or 30 μM E2 alone. Samples omitted if not responsive to the desired pacing frequency*.

## DISCUSSION

To the best of our knowledge, this is the first *in vitro* study to compare the effects of bisphenol chemicals and estradiol on cardiac physiology using a human relevant model. Key findings of our study include: 1) bisphenol chemicals and estradiol alter cardiac physiology acutely and reversibly (within minutes), indicating non-genomic activity that likely occurs before these chemicals are metabolized and excreted. 2) Despite similarities in structure, the potency of estradiol and bisphenol chemicals differ. Across multiple cardiac metrics, an approximate potency hierarchy includes E2 ≥ BPA ≥ BPF >> BPS. 3) BPA and E2 cause similar effects on cardiomyocytes (e.g. FPD and APD shortening, smaller depolarizing spike amplitude), but the underlying mechanisms are not identical. 4) Although there were a few exceptions (e.g., APD shortening), perturbations in cardiac physiology were largely observed at elevated micromolar concentrations. This finding highlights the fact that hiPSC-CM are a useful screening tool, but cellular models do not fully replicate the intact heart and/or cardiovascular system – and key findings should be validated with a higher-order cardiac model (e.g., intact electrical conduction system, atrial versus ventricular chamber differences, contractile function against a mechanical load).

Our findings highlight the acute, non-genomic activity of bisphenol chemicals – which aligns with previous experimental studies. Both BPA and E2 have been shown to interfere with intracellular calcium handling in isolated ventricular cardiomyocytes and hiPSC-CM within a short timeframe (2-15 minutes)(Hyun et al. 2021; Jiang et al. 1992; Ramadan et al. 2018; Yan et al. 2011). Potential mechanisms underlying the effects on calcium handling and contractility include the phosphorylation of key calcium handling proteins (Gao et al. 2013; Liang et al. 2014), and/or the inhibition of inward calcium current (Deutschmann et al. 2013; Hyun et al. 2021; Jiang et al. 1992; Liang et al. 2014; Meyer et al. 1998; Prudencio et al. 2021). Although the reported potency of BPA on L-type calcium channels does vary across models, ranging from ∼16.5% at 1 nM BPA in hiPSC-CM, ∼12% at 10 nM in rat ventricular cardiomyocytes, and ∼10% at 5 μM BPA in transfected cell lines (Liang et al. 2014; Ma et al. 2023b; Prudencio et al. 2021). As expected with calcium channel inhibitors, we found that both BPA and E2 hasten repolarization by shortening the APD and FPD in hiPSC-CM in a dose-dependent manner, which aligns with previous work (Hyun et al. 2021; Jiang et al. 1992; Prudencio et al. 2021). Of interest, at least one study suggests that BPA exposure can delay repolarization through the inhibition of hERG channels at low nanomolar concentrations, before the onset of calcium channel inhibition at higher concentrations (Ma et al. 2023a). Nevertheless, the effects of E2 and BPA on cardiac electrophysiology and calcium handling have been shown to occur quickly and reversibly.

In the presented study, we observed similar perturbations in hiPSC-CM physiology upon exposure to either BPA or E2; yet, our pharmacological studies indicate that these two chemicals have a few distinct features. First, pretreatment with either an L-type calcium channel antagonist or ERα agonist exaggerated the effects of BPA on beating rate and FPD – but a similar effect was not observed with E2. Second, pretreatment with an ERα antagonist reduced (albeit only slightly) the effects of BPA on beating rate and FPD – but this was not observed with E2. Notably, a link between BPA cardiotoxicity and estrogen receptor signaling has previously been documented, whereby the effects of BPA were abolished in an ERβ knockout mouse or following treatment with an ERβ blocker (Belcher et al. 2011; Gao et al. 2013; Liang et al. 2014; Yan et al. 2011). Collectively, these studies highlight the complexity of BPA activity in the heart, which can be influenced by multiple factors including its interaction with cardiac ion channels, as well as estrogen receptor signaling.

Consistent with our earlier work (Prudencio et al. 2021), this study confirmed that the potency of E2 and bisphenol chemicals differ markedly. We previously showed that BPA is a more potent inhibitor of cardiac ion channel currents (*I*_Na_, *I*_Ca,L_, *I*_Kr_), as compared to either BPF or BPS. Pharmacological studies suggest that bisphenol chemicals physically interact with the extracellular component of voltage-gated calcium channels – and this interaction is influenced by the angular orientation of each bisphenol compound (calcium channel inhibition: BPA > BPF > BPS) (Deutschmann et al. 2013). Further, a recent study comparing 14 different bisphenol chemicals using aortic ring preparations reported that BPS had negligible biological effects (Tvrdý et al. 2023). In agreement, we report here that BPS had negligible effects on hiPSC-CM beating rate, FPD, APD, or CaD – suggesting it may offer a superior cardiac safety profile as compared to BPA. However, future work is still required to extensively study these replacement chemicals, as others groups have reported that BPS can interfere with calcium handling and contractility in (rodent) ventricular cardiomyocytes or intact heart preparations (Ferguson et al. 2019; Gao et al. 2015). This discrepancy may be attributed to the species differences and the experimental model used in each study. As examples, rodent ventricular cardiomyocytes have a shorter action potential duration and rely more heavily on the sarcoendoplasmic reticulum calcium ATPase for calcium removal compared to human cells (Bers 2008; Ripplinger et al. 2022); yet, hiPSC-CM do not fully replicate adult human ventricular myocytes given their automaticity and immature calcium handling machinery (Karbassi et al. 2020; Salameh et al. 2023). Nevertheless, comparative studies have demonstrated excellent correlation between FPDc measurements in hiPSC-CM and clinical QTc measurements in response to pharmacological agents – supporting the use of hiPSC-CM as a translational model and screening tool for toxicological assessments (Blinova et al. 2017).

## CONCLUSION AND LIMITATIONS

Epidemiological studies suggest an association between BPA exposure and adverse cardiovascular outcomes (Bao et al. 2020; Cooper and Posnack 2022; Moon et al. 2021), yet there remains a paucity of experimental evidence for the source of bisphenol-induced effects. This study builds upon our previously findings (intact rodent heart model (Prudencio et al. 2021)) – by using hiPSC-CM, which have been regarded as a new alternative approach methodology with high translational potential (Daley et al. 2023). In the current study, we present novel information about the acute effects of bisphenol chemicals (BPA, BPF, BPS) on cardiac electrophysiology – and compare these findings with the most prominent circulating estrogen, 17β-estradiol. Using a broad range of environmentally and clinically relevant doses (7 doses from 0.01-100 μM), we determined that bisphenols have an immediate, non-genomic effect on cardiac electrophysiology and intracellular calcium handling. Observed effects occurred in a dose-dependent manner, with more significant perturbations observed at micromolar concentrations. Of interest, BPS had negligible effects on cardiac physiology, suggesting that it could be a safer replacement chemical. Although these results add important insight to our understanding of bisphenol cardiac toxicity, a few limitations should be considered. First, we utilized hiPSC-CM in our study, which are known to have an immature structural phenotype (e.g., sparse T-tubules, less organized sarcomeres). Nevertheless, these cells have comparable ion channel expression to adult cardiomyocytes – allowing them to serve as a valuable, high-throughput model for cardiotoxicity testing (Karakikes et al. 2015; Koivumäki et al. 2018). Second, it is important to note that data collected from a cellular monolayer is distinct from a coordinated three-dimensional tissue. Although iCell hiPSC-CM are a composition of different cell types, the majority are ventricular cardiomyocytes, therefore these cells may be less sensitive to toxins that target pacemaker or atrial cardiomyocytes. Accordingly, *in vitro* screening studies should be validated using a more complex three-dimensional *ex vivo* or *in vivo* model. The latter also takes into account bisphenol chemical metabolism, which is an important consideration for longer time course studies, as BPA is metabolized and excreted within a few hours (Völkel et al. 2002).

## Abbreviations

APD_80_: action potential duration at 80% repolarization
AV: atrioventricular
BPA: bisphenol A
BPF: bisphenol F
BPS: bisphenol S
CaD_80_: calcium transient duration at 80% reuptake
DMSO: dimethyl sulfoxide
E2: 17β-estradiol
ERα: estrogen receptor alpha
ERβ: estrogen receptor beta
FPDc: field potential duration, Fridericia rate-corrected
hiPSC-CM: human induced pluripotent stem cell-derived cardiomyocytes
IC_50_: 50% inhibitory concentration
MEA: microelectrode array
SR: sarcoplasmic reticulum
SERCA: SR calcium ATPase

## CONFLICT OF INTEREST

The authors declare that the research was conducted in the absence of any commercial or financial relationships that could be construed as a potential conflict of interest.

## AUTHOR CONTRIBUTIONS

BC and SS performed experiments; BC, SS, NGP analyzed data; BC and NGP prepared figures; BC and NGP drafted manuscript; BC and NGP conceived and designed experiments; BC, SS, and NGP approved the manuscript.

## FUNDING

This work was supported by the National Institutes of Health (R01HL139472 and F31HL162549), The Sheikh Zayed Institute for Pediatric Surgical Innovation, and the Children’s Research Institute.

## ACKNOWLEDGEMENTS

The authors gratefully acknowledge Devon Guerrelli, Jenna Pressman, Anysja Roberts, and Sam Allen for technical assistance and helpful scientific discussions related to this project.

## Notes

**Conflict of Interest:** The authors declare no conflicts of interest.

### Competing Interest Statement

The authors have declared no competing interest.

## REFERENCES

Bae S, Hong Y-C. 2015. Exposure to bisphenol A from drinking canned beverages increases blood pressure: randomized crossover trial. Hypertension 65:313–319; doi:10.1161/HYPERTENSIONAHA.114.04261.

Bae S, Kim JH, Lim Y-H, Park HY, Hong Y-C. 2012. Associations of bisphenol A exposure with heart rate variability and blood pressure. Hypertension 60:786–793; doi:10.1161/HYPERTENSIONAHA.112.197715.

Bao W, Liu B, Rong S, Dai SY, Trasande L, Lehmler HJ. 2020. Association Between Bisphenol A Exposure and Risk of All-Cause and Cause-Specific Mortality in US Adults. JAMA Netw Open 3:e2011620; doi:10.1001/jamanetworkopen.2020.11620.

Belcher SM, Chen Y, Yan S, Wang H-SS. 2011. Rapid Estrogen Receptor-Mediated Mechanisms Determine the Sexually Dimorphic Sensitivity of Ventricular Myocytes to 17beta-Estradiol and the Environmental Endocrine Disruptor Bisphenol A. Endocrinology 153:712–720; doi:10.1210/en.2011-1772.

Berger F, Borchard U, Hafner D, Putz I, Weis TM. 1997. Effects of 17beta-estradiol on action potentials and ionic currents in male rat ventricular myocytes. Naunyn Schmiedebergs Arch Pharmacol 356: 788–796.

Bers DM. 2008. Excitation-contraction coupling and cardiac contractile force. Second. Springer.

Blinova K, Stohlman J, Vicente J, Chan D, Johannesen L, Hortigon-Vinagre MP, et al. 2017. Comprehensive Translational Assessment of Human-Induced Pluripotent Stem Cell Derived Cardiomyocytes for Evaluating Drug-Induced Arrhythmias. Toxicological Sciences 155:234–247; doi:10.1093/toxsci/kfw200.

Bousoumah R, Leso V, Iavicoli I, Huuskonen P, Viegas S, Porras SP, et al. 2021. Biomonitoring of occupational exposure to bisphenol A, bisphenol S and bisphenol F: A systematic review. Science of The Total Environment 783:146905; doi:10.1016/j.scitotenv.2021.146905.

Calafat AM, Kuklenyik Z, Reidy JA, Caudill SP, Ekong J, Needham LL. 2005. Urinary concentrations of bisphenol A and 4-Nonylphenol in a human reference population. Environ Health Perspect 113:391–395; doi:10.1289/ehp.7534.

Calafat AM, Ye X, Wong LY, Reidy JA, Needham LL. 2008. Exposure of the U.S. population to Bisphenol A and 4-tertiary-octylphenol: 2003-2004. Environ Health Perspect 116:39–44; doi:10.1289/ehp.10753.

Chen IY, Matsa E, Wu JC. 2016. Induced pluripotent stem cells: at the heart of cardiovascular precision medicine. Nature Publishing Group 13; doi:10.1038/nrcardio.2016.36.

Chen S, Tao Y, Wang P, Li D, Shen R, Fu G, et al. 2023. Association of urinary bisphenol A with cardiovascular and all-cause mortality: National Health and Nutrition Examination Survey (NHANES) 2003-2016. Environ Sci Pollut Res Int 30; doi:10.1007/S11356-023-25924-7.

Cooper BL, Gloschat C, Swift LM, Prudencio T, McCullough D, Jaimes R 3rd, et al. 2021. KairoSight: Open-Source Software for the Analysis of Cardiac Optical Data Collected From Multiple Species. 1820; doi:10.3389/FPHYS.2021.752940.

Cooper BL, Posnack NG. 2022. Characteristics of Bisphenol Cardiotoxicity: Impaired Excitability, Contractility, and Relaxation. Cardiovasc Toxicol 22:273–280; doi:10.1007/S12012-022-09719-9.

Daley MC, Mende U, Choi BR, McMullen PD, Coulombe KLK. 2023. Beyond pharmaceuticals: Fit-for-purpose new approach methodologies for environmental cardiotoxicity testing. ALTEX - Alternatives to animal experimentation 40:103–116; doi:10.14573/ALTEX.2109131.

Delfosse V, Grimaldi M, Pons JL, Boulahtouf A, Le Maire A, Cavailles V, et al. 2012. Structural and mechanistic insights into bisphenols action provide guidelines for risk assessment and discovery of bisphenol A substitutes. Proc Natl Acad Sci U S A 109:14930–14935; doi:10.1073/PNAS.1203574109.

Deutschmann A, Hans M, Meyer R, Häberlein H, Swandulla D. 2013. Bisphenol A inhibits voltage-activated Ca(2+) channels in vitro: mechanisms and structural requirements. Mol Pharmacol 83:501–511; doi:10.1124/mol.112.081372.

Duty SM, Mendonca K, Hauser R, Calafat AM, Ye X, Meeker JD, et al. 2013. Potential sources of bisphenol a in the neonatal intensive care unit. Pediatrics 131:483–489; doi:10.1542/peds.2012-1380.

Edwards AG, Louch WE. 2017. Species-Dependent Mechanisms of Cardiac Arrhythmia: A Cellular Focus. Clin Med Insights Cardiol 11; doi:10.1177/1179546816686061.

Feiteiro J, Mariana M, Glória S, Cairrao E. 2018. Inhibition of L-type calcium channels by Bisphenol A in rat aorta smooth muscle. Journal of Toxicological Sciences 43:579–586; doi:10.2131/jts.43.579.

Ferguson M, Lorenzen-Schmidt I, Pyle WG. 2019. Bisphenol S rapidly depresses heart function through estrogen receptor-β and decreases phospholamban phosphorylation in a sex-dependent manner. Sci Rep 9:15948; doi:10.1038/s41598-019-52350-y.

Gao X, Liang Q, Chen Y, Wang H-SS. 2013. Molecular mechanisms underlying the rapid arrhythmogenic action of bisphenol A in female rat hearts. Endocrinology 154:4607–4617; doi:10.1210/en.2013-1737.

Gao X, Ma J, Chen Y, Wang H-S. 2015. Rapid responses and mechanism of action for low-dose bisphenol S on ex vivo rat hearts and isolated myocytes: evidence of female-specific proarrhythmic effects. Environ Health Perspect 123:571–8; doi:10.1289/ehp.1408679.

Gintant G, Fermini B, Stockbridge N, Strauss D. 2017. The Evolving Roles of Human iPSC-Derived Cardiomyocytes in Drug Safety and Discovery. Cell Stem Cell 21:14–17; doi:10.1016/j.stem.2017.06.005.

Hamilton S, Yatani A, Brush K, A S, AM B. 1987. A comparison between the binding and electrophysiological effects of dihydropyridines on cardiac membranes - PubMed. Mol Pharmacol 31: 221–31.

Hayes HB, Nicolini AM, Arrowood CA, Chvatal SA, Wolfson DW, Cho HC, et al. 2019. Novel method for action potential measurements from intact cardiac monolayers with multiwell microelectrode array technology. Sci Rep 9:11893; doi:10.1038/s41598-019-48174-5.

Hyun S-A, Lee CY, Ko MY, Chon S-H, Kim Y-J, Seo J-W, et al. 2021. Cardiac toxicity from bisphenol A exposure in human-induced pluripotent stem cell-derived cardiomyocytes. Toxicol Appl Pharmacol 428:115696; doi:10.1016/j.taap.2021.115696.

Iribarne-Durán LM, Artacho-Cordón F, Peña-Caballero M, Molina-Molina JM, Jiménez-Díaz I, Vela-Soria F, et al. 2019. Presence of bisphenol a and parabens in a neonatal intensive care unit: An exploratory study of potential sources of exposure. Environ Health Perspect 127; doi:10.1289/EHP5564.

Jaimes R, Walton RD, Pasdois PPLCP, Bernus O, Efimov IR, Kay MW. 2016. A technical review of optical mapping of intracellular calcium within myocardial tissue. American Journal of Physiology-Heart and Circulatory Physiology 310:H1388–401; doi:10.1152/ajpheart.00665.2015.

Jiang C, Poole-Wilson PA, Sarrel PM, Mochizuki S, Collins P, MacLeod KT. 1992. Effect of 17 beta-oestradiol on contraction, Ca2+ current and intracellular free Ca2+ in guinea-pig isolated cardiac myocytes. Br J Pharmacol 106:739–745; doi:10.1111/J.1476-5381.1992.TB14403.X.

Karakikes I, Ameen M, Termglinchan V, Wu JC. 2015. Human induced pluripotent stem cell-derived cardiomyocytes: insights into molecular, cellular, and functional phenotypes. Circ Res 117:80–88; doi:10.1161/CIRCRESAHA.117.305365.

Karbassi E, Fenix A, Marchiano S, Muraoka N, Nakamura K, Yang X, et al. 2020. Cardiomyocyte maturation: advances in knowledge and implications for regenerative medicine. Nat Rev Cardiol; doi:10.1038/s41569-019-0331-x.

Koivumäki JT, Naumenko N, Tuomainen T, Takalo J, Oksanen M, Puttonen KA, et al. 2018. Structural immaturity of human iPSC-derived cardiomyocytes: In silico investigation of effects on function and disease modeling. Front Physiol 9:319019; doi:10.3389/FPHYS.2018.00080/BIBTEX.

Kojima H, Takeuchi S, Sanoh S, Okuda K, Kitamura S, Uramaru N, et al. 2019. Profiling of bisphenol A and eight of its analogues on transcriptional activity via human nuclear receptors. Toxicology 413:48–55; doi:10.1016/J.TOX.2018.12.001.

Lehmler HJ, Liu B, Gadogbe M, Bao W. 2018. Exposure to Bisphenol A, Bisphenol F, and Bisphenol S in U.S. Adults and Children: The National Health and Nutrition Examination Survey 2013-2014. ACS Omega 3:6523–6532; doi:10.1021/acsomega.8b00824.

Liang Q, Gao X, Chen Y, Hong K, Wang H-SS. 2014. Cellular mechanism of the nonmonotonic dose response of bisphenol A in rat cardiac myocytes. Environ Health Perspect 122:601–608; doi:10.1289/ehp.1307491.

Lind L, Araujo JA, Barehowsky A, Belcher S, Berridge BR, Chiamvimonvat N, et al. 2021. Key characteristics of cardiovascular Toxicants. Environ Health Perspect 129:95001–95001; doi:10.1289/EHP9321.

Long BJ, Tilghman SL, Yue W, Thiantanawat A, Grigoryev DN, Brodie AMH. 1998. The steroidal antiestrogen ICI 182,780 is an inhibitor of cellular aromatase activity. J Steroid Biochem Mol Biol 67:293–304; doi:10.1016/S0960-0760(98)00122-8.

Ma J, Hong K, Wang H-SS. 2017. Progesterone Protects Against Bisphenol A–Induced Arrhythmias in Female Rat Cardiac Myocytes via Rapid Signaling. Endocrinology 158:778–790; doi:10.1210/en.2016-1702.

Ma J, Niklewski PJ, Wang HS. 2023a. Acute exposure to low-dose bisphenol A delays cardiac repolarization in female canine heart - Implication for proarrhythmic toxicity in large animals. Food Chem Toxicol 172; doi:10.1016/J.FCT.2022.113589.

Ma J, Wang NY, Jagani R, Wang H-S. 2023b. Proarrhythmic toxicity of low dose bisphenol A and its analogs in human iPSC-derived cardiomyocytes and human cardiac organoids through delay of cardiac repolarization. Chemosphere 328:138562; doi:10.1016/J.CHEMOSPHERE.2023.138562.

Melzer D, Rice NE, Lewis C, Henley WE, Galloway TS. 2010. Association of urinary bisphenol a concentration with heart disease: evidence from NHANES 2003/06. PLoS One 5:e8673–e8673; doi:10.1371/journal.pone.0008673.

Meyer R, Linz KW, Surges R, Meinardus S, Vees J, Hoffmann A, et al. 1998. Rapid modulation of L-type calcium current by acutely applied oestrogens in isolated cardiac myocytes from human, guinea-pig and rat. Exp Physiol 83:305–321; doi:10.1113/EXPPHYSIOL.1998.SP004115.

Michaela P, Mária K, Silvia H, Ľubica L, L’ubica L. 2014. Bisphenol A Differently Inhibits CaV3.1, Ca V3.2 and Ca V3.3 Calcium Channels. Naunyn Schmiedebergs Archives of Pharmacology 387; doi:10.1007/s00210-013-0932-6.

Moon S, Yu SH, Lee CB, Park YJ, Yoo HJ, Kim DS. 2021. Effects of bisphenol A on cardiovascular disease: An epidemiological study using National Health and Nutrition Examination Survey 2003– 2016 and meta-analysis. Science of the Total Environment 763:142941; doi:10.1016/j.scitotenv.2020.142941.

O’Reilly AO, Eberhardt E, Weidner C, Alzheimer C, Wallace BA, Lampert A. 2012. Bisphenol a binds to the local anesthetic receptor site to block the human cardiac sodium channel. PLoS One 7:e41667–e41667; doi:10.1371/journal.pone.0041667.

Pant J, Ranjan P, Deshpande SB. 2011. Bisphenol A decreases atrial contractility involving NO-dependent G-cyclase signaling pathway. Journal of Applied Toxicology 31:698–702; doi:10.1002/JAT.1647.

Parish ST, Aschner M, Casey W, Corvaro M, Embry MR, Fitzpatrick S, et al. 2020. An evaluation framework for new approach methodologies (NAMs) for human health safety assessment. Regul Toxicol Pharmacol 112; doi:10.1016/J.YRTPH.2020.104592.

Paterni I, Granchi C, Katzenellenbogen JA, Minutolo F. 2014. Estrogen Receptors Alpha (ERα) and Beta (ERβ): Subtype-Selective Ligands and Clinical Potential. Steroids 0:13; doi:10.1016/J.STEROIDS.2014.06.012.

Posnack NG, Brooks D, Chandra A, Jaimes R, Sarvazyan N, Kay MW. 2015. Physiological response of cardiac tissue to Bisphenol A: alterations in ventricular pressure and contractility. Am J Physiol Heart Circ Physiol 309:H267–H275; doi:10.1152/ajpheart.00272.2015.

Posnack NG, Jaimes III R, Asfour H, Swift LM, Wengrowski AM, Sarvazyan N, et al. 2014. Bisphenol A Exposure and Cardiac Electrical Conduction in Excised Rat Hearts. Environ Health Perspect 122:384–90; doi:10.1289/ehp.1206157.

Prudencio TM, Swift LM, Guerrelli D, Cooper B, Reilly M, Ciccarelli N, et al. 2021. Bisphenol S and Bisphenol F Are Less Disruptive to Cardiac Electrophysiology, as Compared With Bisphenol A. Toxicological Sciences 183:214–226; doi:10.1093/toxsci/kfab083.

Ramadan M, Cooper B, Posnack NG. 2020. Bisphenols and phthalates: Plastic chemical exposures can contribute to adverse cardiovascular health outcomes. Birth Defects Res; doi:10.1002/bdr2.1752.

Ramadan M, Sherman M, Jaimes R, Chaluvadi A, Swift L, Posnack NGNG, et al. 2018. Disruption of neonatal cardiomyocyte physiology following exposure to bisphenol-a. Sci Rep 8:7356; doi:10.1038/s41598-018-25719-8.

Ripplinger CM, Glukhov A V., Kay MW, Boukens BJ, Chiamvimonvat N, Delisle BP, et al. 2022. Guidelines for assessment of cardiac electrophysiology and arrhythmias in small animals. Am J Physiol Heart Circ Physiol 323: H1137–H1166.

Salameh S, Ogueri V, Posnack NG. 2023. Adapting to a new environment: postnatal maturation of the human cardiomyocyte. J Physiol; doi:10.1113/JP283792.

Shankar A, Teppala S. 2012. Urinary bisphenol A and hypertension in a multiethnic sample of US adults. J Environ Public Health 2012:481641; doi:10.1155/2012/481641.

Sinnecker D, Goedel A, Laugwitz K-L, Moretti A. 2013. Induced Pluripotent Stem Cell-Derived Cardiomyocytes: A Versatile Tool for Arrhythmia Research. Circ Res 112:961–968; doi:10.1161/CIRCRESAHA.112.268623.

Trasande L. 2017. Exploring regrettable substitution: replacements for bisphenol A. Lancet Planet Health 1:e88–e89; doi:10.1016/S2542-5196(17)30046-3.

Tvrdý V, Dias P, Nejmanová I, Carazo A, Jirkovský E, Pourová J, et al. 2023. The effects of bisphenols on the cardiovascular system ex vivo and in vivo. Chemosphere 313; doi:10.1016/J.CHEMOSPHERE.2022.137565.

Vandenberg LN, Chahoud I, Heindel JJ, Padmanabhan V, Paumgartten FJR, Schoenfelder G. 2010. Urinary, circulating, and tissue biomonitoring studies indicate widespread exposure to bisphenol A. Environ Health Perspect 118:1055–70; doi:10.1289/ehp.0901716.

Vandenberg LN, Hauser R, Marcus M, Olea N, Welshons W V. 2007. Human exposure to bisphenol A (BPA). Reproductive Toxicology 24:139–177; doi:10.1016/j.reprotox.2007.07.010.

Völkel W, Colnot T, Csanády GA, Filser JG, Dekant W. 2002. Metabolism and kinetics of bisphenol a in humans at low doses following oral administration. Chem Res Toxicol 15:1281–1287; doi:10.1021/TX025548T.

Wang Q, Cao J, Zhu Q, Luan C, Chen X, Yi X, et al. 2011. Inhibition of voltage-gated sodium channels by bisphenol A in mouse dorsal root ganglion neurons. Brain Res 1378:1–8; doi:10.1016/j.brainres.2011.01.022.

Wang R, Fei Q, Liu S, Weng X, Liang H, Wu Y, et al. 2022. The bisphenol F and bisphenol S and cardiovascular disease: results from NHANES 2013–2016. Environ Sci Eur 34:1–10; doi:10.1186/S12302-021-00586-9/TABLES/3.

Yan S, Chen Y, Dong M, Song W, Belcher SM, Wang HS. 2011. Bisphenol A and 17β-estradiol promote arrhythmia in the female heart via alteration of calcium handling. PLoS One 6:e25455; doi:10.1371/journal.pone.0025455.

Yan S, Song W, Chen Y, Hong K, Rubinstein J, Wang H-SS. 2013. Low-dose bisphenol A and estrogen increase ventricular arrhythmias following ischemia-reperfusion in female rat hearts. Food Chem Toxicol 56:75–80; doi:10.1016/j.fct.2013.02.011.

Ye X, Wong LY, Kramer J, Zhou X, Jia T, Calafat AM. 2015. Urinary Concentrations of Bisphenol A and Three Other Bisphenols in Convenience Samples of U.S. Adults during 2000-2014. Environ Sci Technol 49:11834–11839; doi:10.1021/acs.est.5b02135.

Yu X, Xue J, Yao H, Wu Q, Venkatesan AK, Halden RU, et al. 2015. Occurrence and estrogenic potency of eight bisphenol analogs in sewage sludge from the U.S. EPA targeted national sewage sludge survey. J Hazard Mater 299:733–739; doi:10.1016/J.JHAZMAT.2015.07.012.

Zhou HB, Carlson KE, Stossi F, Katzenellenbogen BS, Katzenellenbogen JA. 2009. Analogs of methyl-piperidinopyrazole (MPP): Antiestrogens with estrogen receptor α selective activity. Bioorg Med Chem Lett 19:108–110; doi:10.1016/J.BMCL.2008.11.006.

